# Dopaminergic Gene Dosage in Autism versus Developmental Delay: From Complex Networks to Machine Learning approaches

**DOI:** 10.1101/2020.04.28.065987

**Authors:** André Santos, Francisco Caramelo, Joana Barbosa de Melo, Miguel Castelo-Branco

## Abstract

The neural basis of behavioural changes in Autism Spectrum Disorders (ASD) remains a controversial issue. One factor contributing to this challenge is the phenotypic heterogeneity observed in ASD, which suggests that several different system disruptions may contribute to diverse patterns of impairment between and within study samples. Here, we took a retrospective approach, using SFARI data to study ASD by focusing on participants with genetic imbalances targeting the dopaminergic system. Using complex network analysis, we investigated the relations between participants, Gene Ontology (GO) and gene dosage related to dopaminergic neurotransmission from a polygenic point of view. We converted network analysis into a machine learning binary classification problem to differentiate ASD diagnosed participants from DD (developmental delay) diagnosed participants. Using 1846 participants to train a Random Forest algorithm, our best classifier achieved on average a diagnosis predicting accuracy of 85.18% (sd 1.11%) on a test sample of 790 participants using gene dosage features. In addition, we observed that if the classifier uses GO features it was also able to infer a correct response based on the previous examples because it is tied to a set of biological process, molecular functions and cellular components relevant to the problem. This yields a less variable and more compact set of features when comparing with gene dosage classifiers. Other facets of knowledge-based systems approaches addressing ASD through network analysis and machine learning, providing an interesting avenue of research for the future, are presented through the study.

**Lay Summary:** There are important issues in the differential diagnosis of Autism Spectrum Disorders. Gene dosage effects may be important in this context. In this work, we studied genetic alterations related to dopamine processes that could impact brain development and function of 2636 participants. On average, from a genetic sample we were able to correctly separate autism from developmental delay with an accuracy of 85%.

## 1. Introduction

Autism Spectrum Disorder(ASD) is a neurodevelopmental disorder characterized by social and communication impairments and repetitive behaviours, which biological basis is still poorly understood (Sanders, 2015). In the last years, many genetic variations have been associated to ASD, however, in most cases these patterns were only verified in small size samples leading to a still unclear scenario (Ecker et al., 2010; Lombardo, Lai, & Baron-Cohen, 2019; Masi, DeMayo, Glozier, & Guastella, 2017; Tang et al., 2020) The large range of different individual manifestations and still unknown gene-environment interactions further compounds this problem (Amaral, Schumann, & Nordahl, 2008; Sharma, Gonda, & Tarazi, 2018). Research on putative causal aspects that conduct to the different manifestations of ASD remains a priority. The role of dopaminergic neurotransmission and its involvement in the causal pathway of ASD deserves attention in this regard (Ayano, 2016; Greene, Walsh, Mosner, & Dichter, 2018; Paval, 2017). For example, disruptions in the nigrostriatal pathway, classically linked to motor functions, has been proposed as a potential contributing cause of some motor repetitive behaviours in ASD whereas disturbances in the mesocorticolimbic pathway, involved in reward and cognitive related functions, may lead to social cognitive an affective impairments,(Fernández, Mollinedo-Gajate, & Peñagarikano, 2018; Supekar et al., 2018).

Machine Learning (ML) approaches have been proved useful in many health-related classification tasks. One of the strongest example is its employment on the analysis of radiological images to detect diseases (Zhang & Sejdić, 2019). With respect to ASD, ML classification has been applied in several studies to predict diagnosis, however with sparse prediction results. A 2019 review on the subject (Wolfers et al., 2019), observed a discrepancy in the diagnostic prediction accuracy between 60% and 98% across 57 different studies. The reason given for such disparity was attributed to several aspects, for example, bias in the sample, small sample sizes, usage of different validation methods, the heterogeneity of ASD, and the quality of the data. Another key factor that could be observed in this review is related to the scientific field where the classification tasks were applied. We detected that of the 57 studies, only one focus on genetic features to predict ASD diagnosis. On this particular study (Ghafouri-Fard et al., 2019), 487 ASD patients and 455 healthy individuals were used to build a ML classifier which relied on single nucleotide polymorphism (SNP) data to predict the diagnosis. The accuracy, sensitivity and specificity achieved by their model were 73.67%, 82.75% and 63.95%, respectively. Here, we decided to focus on the distinction between ASD and DD, as an important intermediate step, and also relevant for differential diagnosis. We will present a ML approach to predict diagnosis with gene Copy Number Variations (CNV), aiming to address the issues present earlier and, to increase the number and variety of studies using genetic data to predict ASD diagnosis.

In order to contribute to the discussion of the impact of the dopaminergic system in ASD, we studied several dopaminergic features coded in the gene CNVs of ASD carriers (Vicari et al., 2019). A CNV is a type of genetic alteration affecting several nucleotides in a chromosomic region. This type of alteration can occur in a gene region. Here, we focus on gene CNVs where genes were in duplicated or deleted chromosomic regions. Firstly, we used QuickGO (Binns et al., 2009) to identify a set of Gene Ontology (GO) (Ashburner et al., 2000; Carbon et al., 2019) terms related to the dopaminergic system. Next, from SFARI Gene CNV Module (Fischbach & Lord, 2010)(sfari.org/resource/sfari-gene/) we selected participants having CNVs matching genes of interest based on the previous defined GO terms. To address and study the genetic variance and frequency, heterogeneity, shown in ASD we used complex network analysis (Conte et al., 2019) This approach also allowed to model the data accordingly to a functional polygenic view (Persico & Napolioni, 2013; Visscher et al., 2017), In the end, we transformed the networks into information suited for a ML classification problem. We used Random Decisions Forests (Ho, 1995) to predict the participants’ differential diagnosis when presented with dopamine related features extracted from gene alterations observed in participants.

## 2. Materials and Methods

With QuickGO API we created a set of 110 GO terms related to dopaminergic neurotransmission aspects that were used to identify a set of 125 genes from CNVs dataset curated by the SFARI Gene CNV module. The genes within each CNV regions were identified using the BioMart API (S. Durinck et al., 2005; Steffen Durinck, Spellman, Birney, & Huber, 2009). For this study, were admitted participants of SFARI Gene CNV module with duplications or deletions in the genes identified previously. Participants with diagnostic comorbidities were excluded. Overall, were selected 1318 participants with diagnosis of ASD and 1327 participants with diagnosis of developmental delay (DD).

We continue our approach by building a network of participants and gene alterations (Goh et al., 2007). In this network each participant is linked to a gene if a duplication or a deletion is present. Then, a network of GO terms and participants was built by replacing each gene with the associated GO terms. Lastly, we joined this information into a single network, consisting of links between genes, GO terms and participants. All networks were built using NetworkX package (Hagberg, Schult, & Swart, 2008) and layout with Gephi software (Bastian, Heymann, & Jacomy, 2009) to show a spatial arrangement of the nodes and their links. Node degree was used to size nodes and colours were used to mark different types of nodes. The Fruchterman Reingold (FR) algorithm (Fruchterman & Reingold, 2017) organized the nodes using a gravity approach where the higher the degree of a node the stronger is the force by which it attracts the linked nodes and pushes the unlinked ones. In order to enable visual comparisons, the parameters used on the FR algorithm, namely the area used to display the nodes, the gravity force and the speed at which changes occurred until the stabilization of the algorithm, were set equal for all networks

From networks topology were extracted centrality measures (Bollobás, 2001; M. Newman, 2018), such as, the number of nodes, the number of edges (links), the average degree and the network diameter. In graph theory, the degree of a node identifies the number of connections/links that a node has, whereas the degree distribution, gives the probability of find a node in a network for a given degree. The degree distribution, of each network was compared to the Poisson curve and the Power Law curve. The Poisson curve represents a distribution of the node degree, where most of the nodes have nearly the same degree with small deviations from the average, this type of curve has a bell-like shape. In this scenario, it is unlikely an existence of a node with higher degree than the average of the network degree. Such a node will be an outlier. Contrarily, the Power Law curve predicts the existence of fewer nodes with higher degree named hubs (outliers in a universe of nodes characterized mainly by lower degree nodes). This approach helped to understand if the networks were closer to the Random Network model, described by the Poisson curve, or to the Scale-Free Network model, described by the Power Law curve (Barabási & Albert, 1999). Lastly, the nodes with major degree of each network were reported as well as the average degree of their neighbourhood. In other words, we characterize hub nodes by their number of links to other nodes and by the average of links their neighbours had, the former helps understanding how many participants were linked to a hub and the latter gives information about the average of participants interactions linked to the hub (Barrat, Barthélemy, Pastor-Satorras, & Vespignani, 2004). These measures help to understand the importance of the hub in the network structure. For example, a hub with average neighbour degree of one represents a hub-and-spoke pattern, meaning that all of its neighbours have in average only one link which could only be a link to the hub. In this scenario, if the hub was removed from the network it will disconnect all the nodes linked to it, breaking the structure of the network.

We transform the features of the created networks in data for a ML classification problem (Hastie, 2017; James, Witten, Hastie, & Tibshirani, 2000). The target classification variable was the differential diagnosis of participants, and to predict it we used features extracted from the network topology. We measured the importance of each non-participant node by dividing the node degree (the number of other nodes attached to it) by the total number of links of the network the node belongs, for each participant if it was linked to a particular node we use its node importance multiplied by the type of the participant gene dosage (Figure 1). For example, in the gene network, each gene was a feature, the value for each feature as given by the node degree over the total number of the network links, which we denoted was network node importance, multiplied by a factor of 1 or −1 representing the participant CVN type, a duplication or deletion, respectively. The gene ontology network was framed in the same way. However, for the last network we constructed the features differently. For a given GO term in the network, its value was given by the importance of the gene node it was linked plus its own node importance, multiplied by the participant CNV type (Figure 1). We obtain three datasets relying on different types of data: 1) gene dosage, 2) GO and 3) combined gene dosage and GO data.

**Figure 1.**
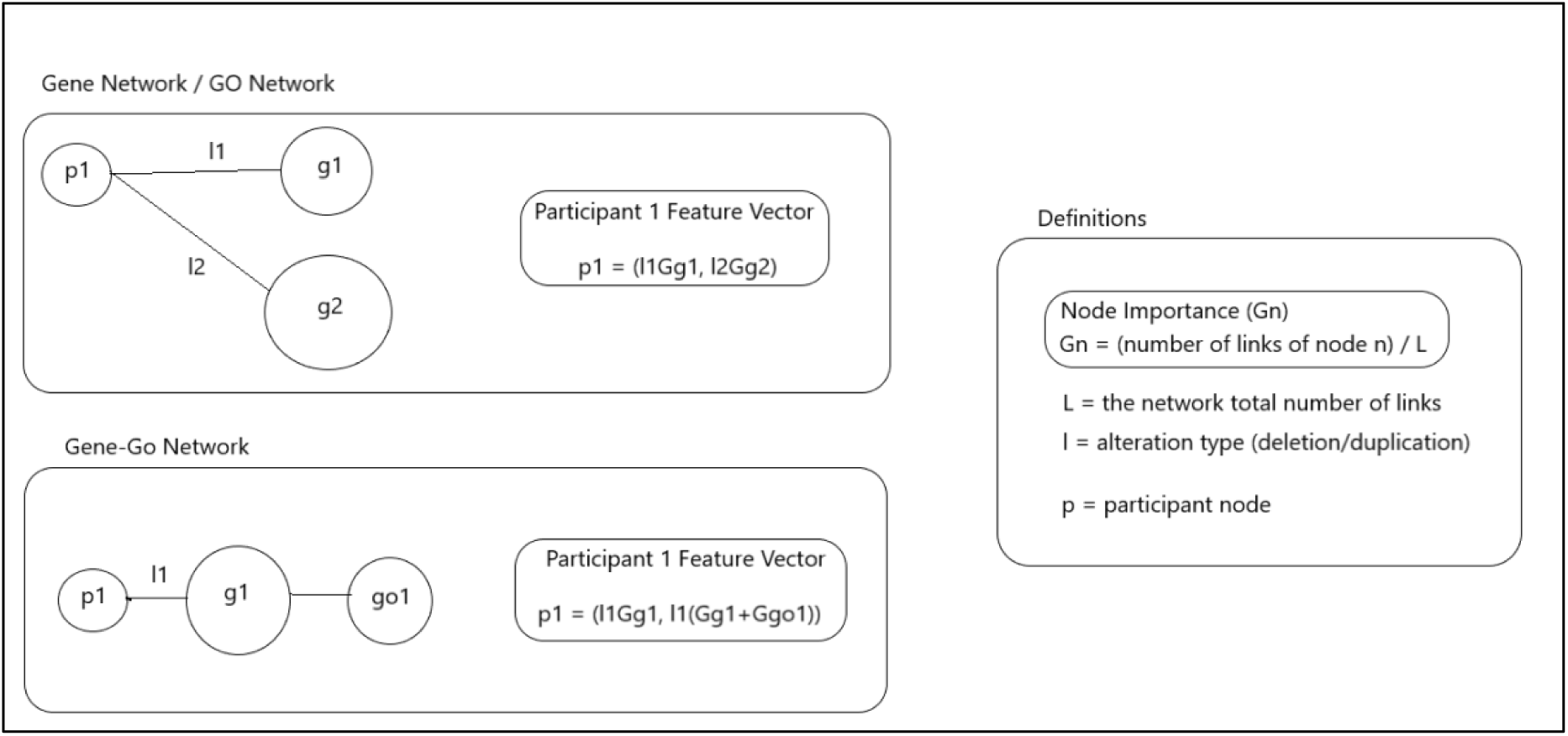
Example of feature extraction from the networks. In Gene Network the vector of features is given by multiplying each gene node importance by the type of alteration (1 for duplications or −1 for deletions) that links a participant to a gene. In GO network the same principle has applied to build the features vector, when a GO term was shared by multiple genes its product was summed. In Gene-Go Network a participant feature vector was built inserting the gene node importance and the GO node importance. However, in this network the GO term was influenced by the importance of the gene node it was linked to. This step allowed to weight differently GO terms shared by multiple genes.

SciKit-Learn package (Varoquaux et al., 2015) was used to train and to test Random Tree classifiers in order to predict the participants diagnosis based on network features and on the participants type of gene dosage, duplication or deletion. We started by random split the participants in a train and a test sets with 70/30 ratio under an equal distribution of classes. In the training set were identified the features with more discriminant power using a wrapper methodology with a threshold of 1e-3 for feature importance (Saeys, Inza, & Larrañaga, 2007) which then were used to train the classifier. The test set was accessed on the previously trained algorithm and the confusion matrix was recorded. This process was repeated 100 times for each type of dataset. At the end, we calculate the mean and the standard deviation of several metrics, such as, the accuracy, the precision, and the recall, from the confusion matrix obtained in each repetition (Sokolova & Lapalme, 2009).

## 3. Results

Data used in this work come from SFARI Gene CNV Module that gathers CNV data related to Autism from several studies. This fact strongly tied our analysis and our results to the scope and the results reported by the original studies to SFARI Gene. In Table I are listed the participants discriminated by diagnosis. About 94% were diagnosed with DD (47.14%) or ASD (46.82%) and the remaining 6% were distributed over other thirteen diagnoses, for example, schizophrenia (1.78%) and intellectual disability (1.67%). Our analysis proceeded with participants having ADS or DD diagnosis due to the differences shown in the frequency of the diagnoses, which led us to focus on this type of differential diagnosis.

**Table I.** Distribution of Participant Diagnosis: Diagnostic and the number of participants (N) identified with duplicated or deleted genes containing information related with dopaminergic aspects on SFARI Gene CNV Module. Only participants with a single diagnostic were considered. * - participants used in this study.

| Diagnostic | N |
| --- | --- |
| Developmental Delay (DD) | 1327* |
| Autism Spectrum Disorder (ASD) | 1318* |
| Schizophrenia | 50 |
| Intellectual Delay (ID) | 41 |
| Middle Cerebral Artery Syndrome (MCA) | 23 |
| Epilepsy | 17 |
| Childhood Apraxia of Speech (CAS) | 13 |
| Polymicrogyria (PMG) | 10 |
| Attention Deficit and Hyperactivity Disorder (ADHD) | 6 |
| Bipolar Disorder | 3 |
| Schizoaffective Disorder | 3 |
| Borderline Personality Disorder (BPD) | 1 |
| Congenital Heart Disease (CHD) | 1 |
| Microcephaly | 1 |
| Angelman Syndrome | 1 |

The methodology used to construct the networks allowed to highlight macroscopic differences between networks (Figure 3, 4 and 5). For example, in the gene dosage network (Figure 3), nodes with larger size were displayed near the limits of the figure and distant from which other, whereas in the GO network (Figure 4) larger nodes appear at the centre of the figure and closer to each other. In the gene network (Figure 3) participants were linked to a single gene or a set of genes, but when we swap the genes nodes by their corresponding GO terms (Figure 4) the links changed, which resulted in an interesting structural alteration of the network showing that many participants share sets of the same dopaminergic aspects. Furthermore, we observed that the percentage of gene nodes (7.44%) in the gene dosage network was higher than the sum of the percentage of the different GO terms nodes (total = 2,73%; biological processes: 1.88%; molecular functions: 0.81%; the cellular component: 0.04%). This feature might be helpful when analysing problems with high genetic variance and low occurrence. Importantly, in this work we were aware that by only looking at one of the networks we were neglecting the information contained in the other. Thus, to overcome this problem, we built the Gene Dosage - GO network (Figure 5) by linking participants, genes and GO terms.

**Figure 2.**
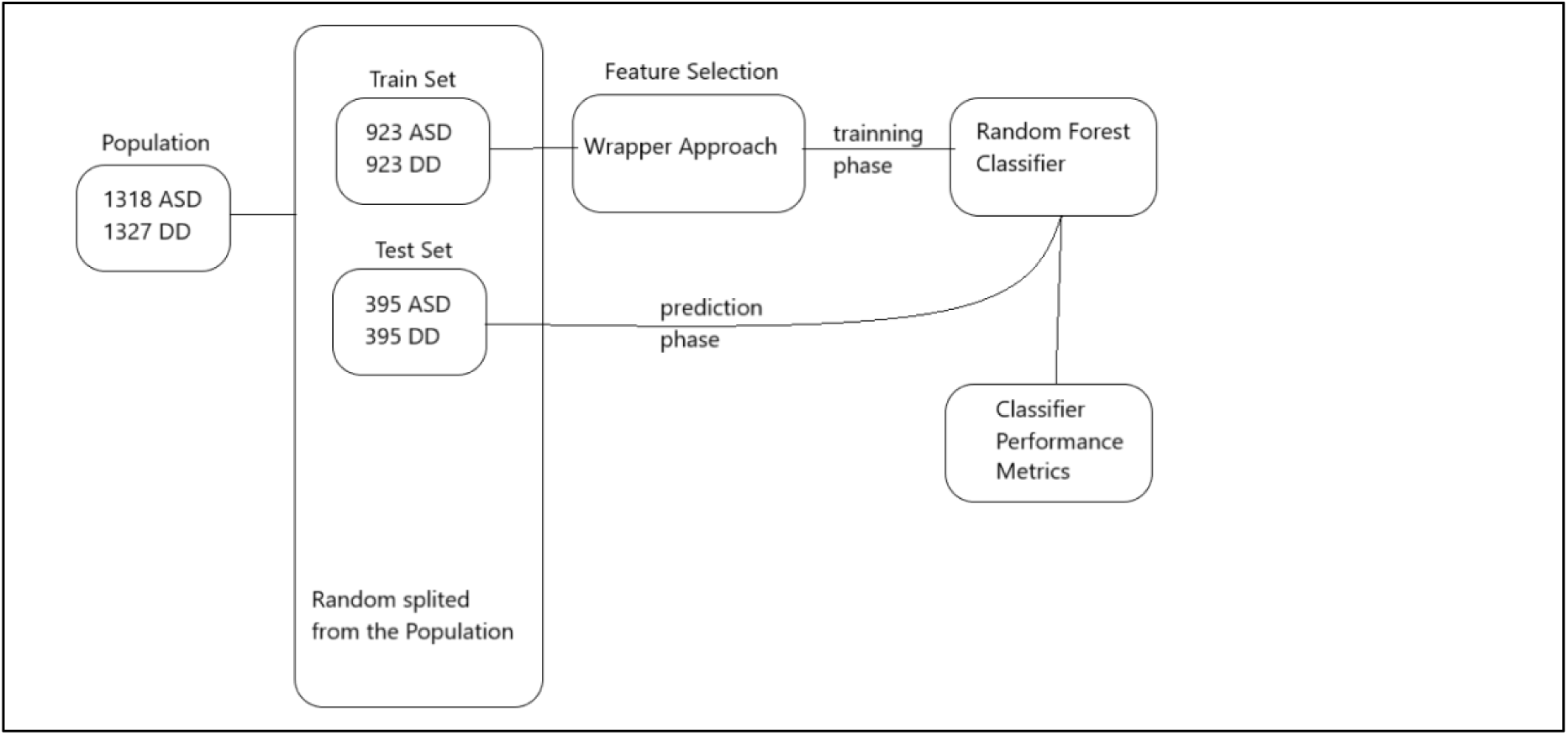
Overview of Machine Learning approach. The population was random split in a training set and a testing set. On the training set was performed a feature reduction using a wrapper approach operating with a Random Forest algorithm. Next, the training set was used to train a Random Forest algorithm. The test set was then accessed by the Random Forest algorithm and the classifier performance was recorded. This process was repeat 100 times.

**Figure 3.**
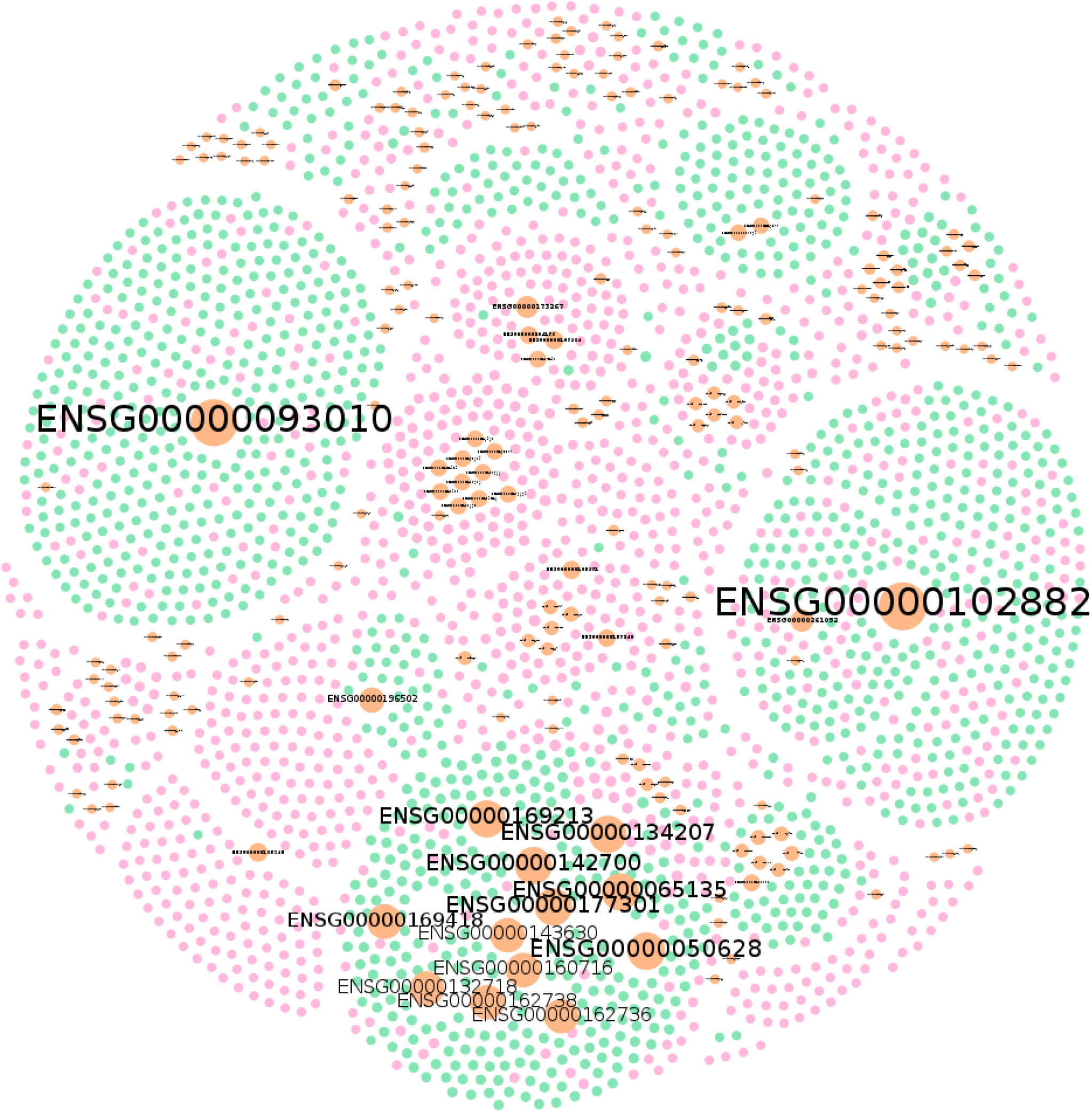
Gene Dosage Network: A network of participants with ASD (pink nodes) or DD (green nodes) diagnosis and their links to duplicated or deleted genes (Orange Nodes). The size of a node reflects its own degree. In this network, ASD nodes represent 45.49% of the total number of the nodes (N on Table II), DD 47.08% and genes 7.44%. Links were omitted for visualization purposes. Produced with Gephi software, using the Fruchterman Reingold Layout (parameters: Area = 10000, Gravity = 5, and Speed = 5)

**Figure 4.**
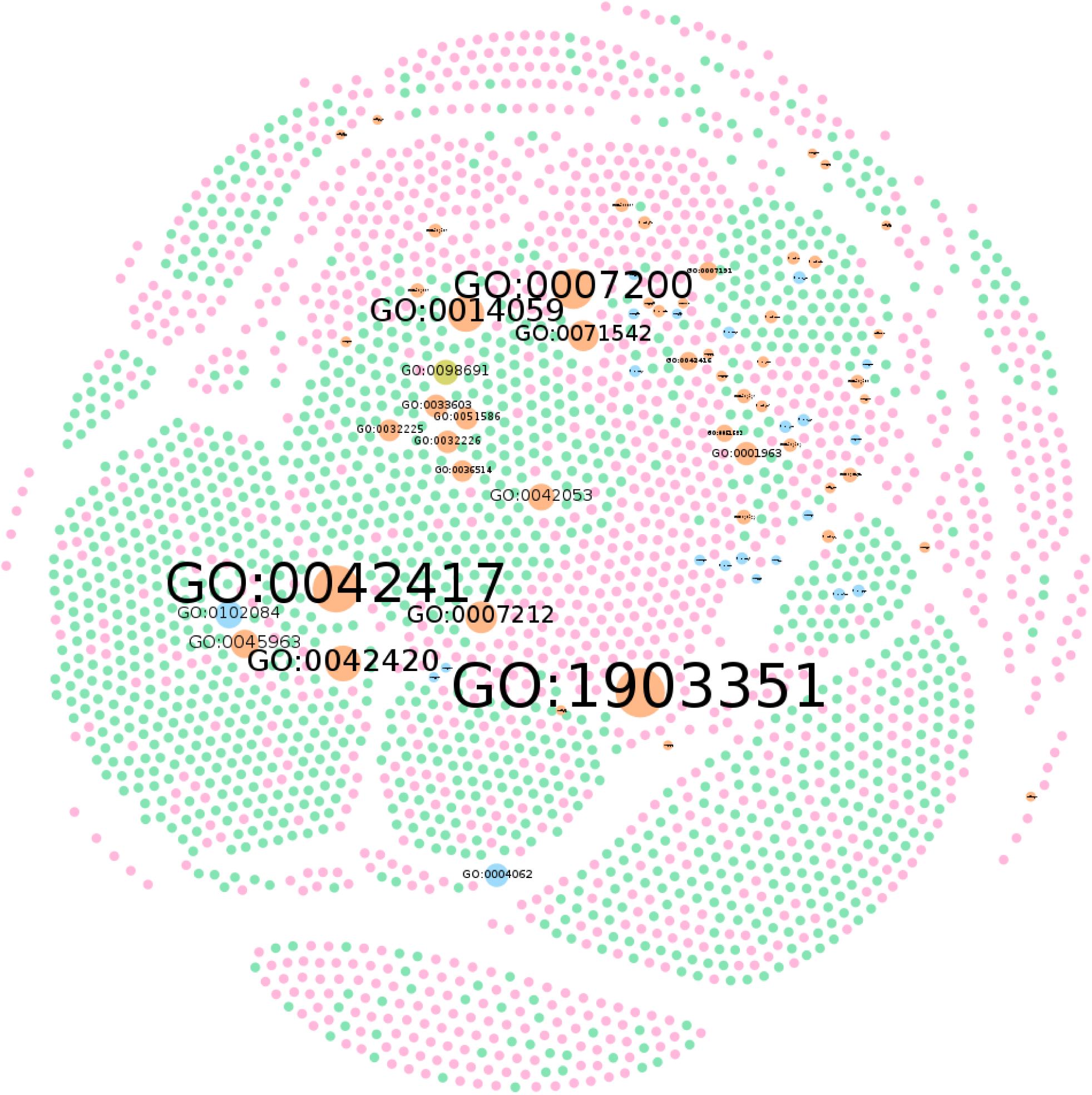
GO network: A network of participants with ASD (pink nodes) or DD (green nodes) diagnosis where the links to their genes were replaced by GO terms: biological processes (orange nodes), molecular functions (blue nodes) and the cellular component (olive node). The size of the node reflects its own degree. In this network, ASD nodes represent 48.43% of the total number of the nodes (N on Table II), DD 48.84%, biological processes 1.88%, molecular functions are 0.81%, and the cellular component 0.04%. Links were omitted for visualization purposes. Produced with Gephi software, using the Fruchterman Reingold Layout (parameters: Area = 10000, Gravity = 5, and Speed = 5)

**Figure 5.**
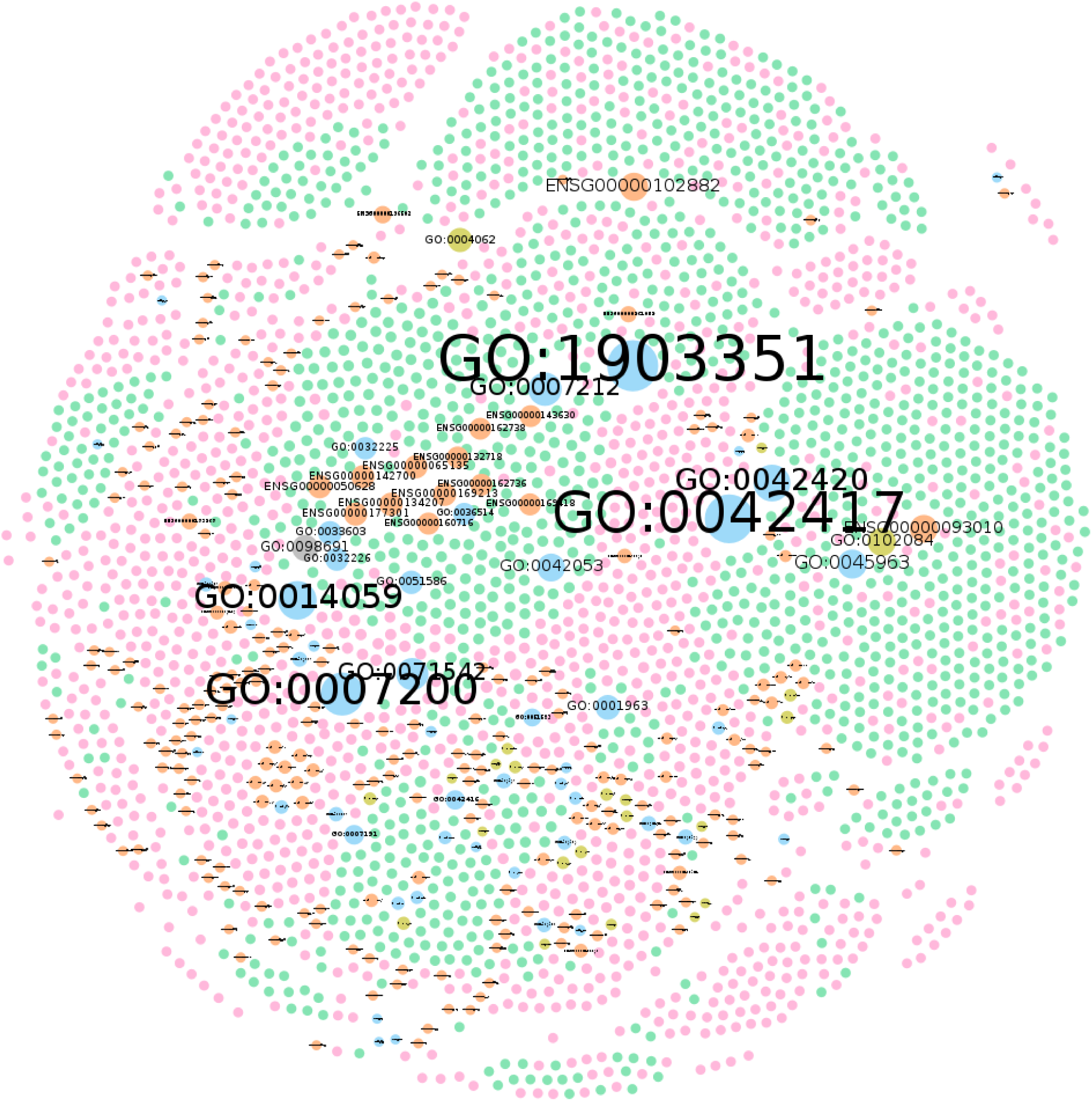
Gene Dosage - GO Network: A network of participants with ASD (pink nodes) or DD (green nodes) diagnosis their links to their genes (orange nodes) and to GO terms: biological processes (blue nodes), molecular functions (olive nodes) and the cellular component (grey node). The size of the node reflects its own degree. In this network, ASD nodes represent 44.65% of the total number of the nodes (N on Table II), DD 44.95%, genes 7.89%, biological processes 1.73%, molecular functions are 0.75%, and the cellular component 0.03%. Links were omitted for visualization purposes. Produced with Gephi software, using the Fruchterman Reingold Layout (parameters: Area = 10000, Gravity = 5, and Speed = 5)

Analyses of centrality measures, degree distribution and hubs helped to confirm the previous macroscopic observations, and to identify new relations between dopaminergic neurotransmission features and the participants differential diagnosis under different contexts. These may be difficult to detect if we relied only on visual inspection analysis. For example, simply counting the number of nodes and links of each network will be extremely difficult. Table II, presents the centrality measures of each network. Looking at the N and L we identify that the gene network was higher N and lower L than the GO network, this decreases the <k> and density properties of the gene network and, comparing to the other two networks we observed that this network is the most poorly connected. Moreover, when we analyse the diameter of the networks, we observe that it is decreased ∼2.3 times from gene network to GO network, this result is in agreement with our initial prediction that in gene network nodes appear more distant than in the GO network. Indeed, by swapping genes by its GO terms we enhanced a property of complex networks known as small worlds (Boldi & Vigna, 2012; de Sola Pool & Kochen, 1978; Travers & Milgram, 1969). In this example, a node can only be at a maximum of a seven nodes distance from another node. By comparing the networks with the Poisson distribution and the Power Law distribution (Faloutsos, Faloutsos, & Faloutsos, 1999; M. E. J. Newman, 2005), Figures 6 and 7, we tried to understand if our networks were closer to a Random Network model, which expects an inexistence of nodes with higher degree than the average network degree, or to a Scale-Free Network model, which expects higher number of nodes with lower degree and existence of fewer hubs, nodes with higher degree (Barabási & Albert, 1999). Here, we observed a long tail behaviour in both networks and assumed the proximity to a Scale-Free model, in other words, in our networks there were nodes with larger degree in a way that they could be labelled as hubs.

**Figure 6.**
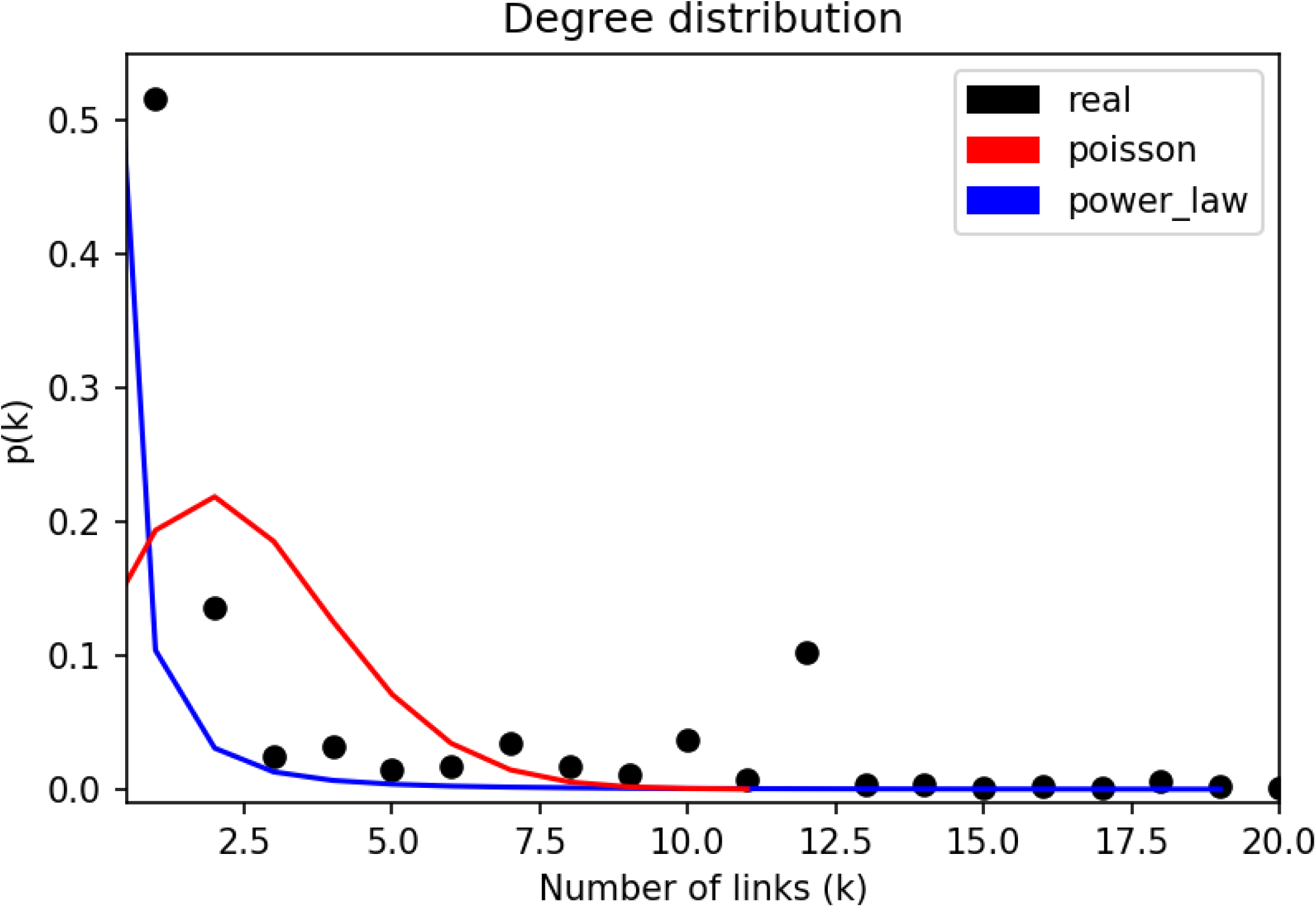
Degree Distribution of the Gene Dosage Network (black dots) and, the Poisson Distribution (red curve) and the Power Law Distribution (blue curve) using the same number of nodes and links of the Gene Dosage Network. On this figure the probability threshold was set to 0.0001 (y axis) and the number of links set to a maximum of 20 (x axis). The maximum number of links was reported on Table III.

**Table II.** Networks Centrality Measures: the total number of nodes (N), the total number links (L), the average degree for unidirectional networks (<k>), the density, the diameter and the radius of each network.

| Network | N | L | $\langle k \rangle$ | Density | Diameter | Radius |
| --- | --- | --- | --- | --- | --- | --- |
| Gene Dosage | 2770 | 9387 | 3.389 | 0.001 | 16 | 8 |
| GO | 2719 | 12669 | 4.659 | 0.002 | 7 | 4 |
| Gene Dosage - GO | 2952 | 22568 | 7.645 | 0.003 | 7 | 4 |

**Figure 7.**
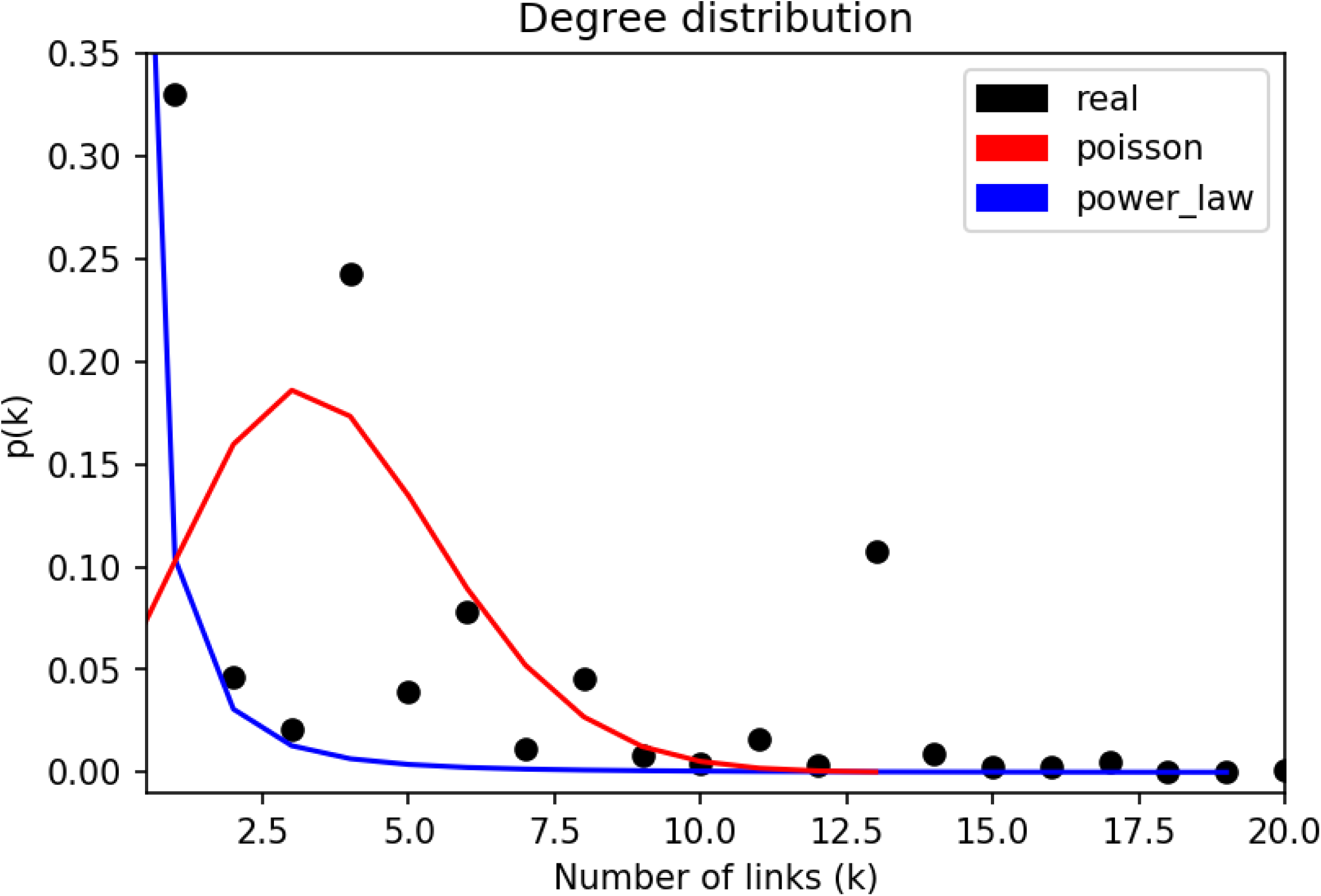
Degree Distribution of GO Network (black dots) and, the Poisson Distribution (red curve) and the Power Law Distribution (blue curve) using the same number of nodes and links of the GO Network. On this figure the probability threshold was set to 0.0001 (y axis) and the number of links set to a maximum of 20 (x axis). The maximum number of links was reported on Table III.

In Table III, were reported the nodes with higher degree and the average degree of its neighbours for each network. For example, in gene network, the major hub (ENSG00000102882) was linked to 463 participants and each one of these participants had on average 1.6 links, meaning that, most of the participants linked to this hub were linked only to it. If this hub was removed from the network, this action will break the connectivity of the network with consequently loss of information. On the other hand, when we look at the last hub (ENSG00000050628) listed for the gene network, we observe that node is linked to 339 participants and an average neighbour degree of 11.8, in that case, each of the participants linked to the hub gene were also linked to other 10.8 genes on average, and removing this hub from the network will not break its connectivity (Albert, Jeong, & Barabási, 2000; Callaway, Newman, Strogatz, & Watts, 2000). Lastly, comparing the hub values of the gene network with the GO network we verify that GO network hubs were linked to more participants, for example, the major hub of GO network had more than twice the links of the major hub of the gene network also, all of the participants linked to hubs in GO network were on average linked to other GO terms where in the gene network it only happens for one hub. Again, Table III supports the visual assumptions made earlier, that in the GO network the GO nodes seemed to be more connected and closer to the centre due to participants being linked to more than one GO term when compared with the gene network. Additionally, from the average neighbour values we observed that GO network in comparison with gene network was more robust, in other words, if one of its hubs were deleted it probably will not break the network structure (Bollobás & Riordan, 2004).

**Table III.** Networks maximum degree nodes (hubs): the node, their name, degree and the average degree of their neighbours of the Gene Dosage and of the GO Networks.

|  |  | Hub | Name | k | Average Neighbours Degree |
| --- | --- | --- | --- | --- | --- |
| Network | Gene Dosage | ENSG00000102882 | MAPK3 | 463 | 1.6 |
|  |  | ENSG00000093010 | COMT | 450 | 1.2 |
|  |  | ENSG00000050628 | PTGER3 | 339 | 11.8 |
|  | GO | GO:1903351 | Cellular response to dopamine | 1029 | 6.3 |
|  |  | GO:0042417 | Dopamine metabolic process | 968 | 7.5 |
|  |  | GO:0007200 | Phospholipase C-activating G protein-coupled receptor signalling pathway | 788 | 8.9 |

In the previous analysis we observed that some nodes had more links than others and, assumed that they had different importance. So, we used it to build features for the classification problem (differential diagnosis between ASD and DD). The results of the ML approach are presented in Table IV and revealed an accuracy at predicting the differential diagnosis of the participants in the test sets of 85.18% (±1.11%), 83.22% (±1.09%) and 85.13% (±1.06%) for Gene Dosage, GO and Gene Dosage - GO datasets, respectively. To understand the types of decision errors that lead to the test accuracy predictions we verified the confusion matrix and inspected both the false positive and false negative predictions. For example, we found that in the Gene Dosage test sets which contained 395 participants diagnosed with ASD and 395 diagnosed with DD on average, 25 (sd 5) ASD participants were misclassified with DD diagnosis and 92(sd 8) DD participants were miss classified with ASD. This pattern is present in the other datasets used in the ML approach and influenced the recall and the precision scores and ultimately the F1 score obtained. Overall, the classifiers ability to identify participants with ASD diagnosis was higher than the ability to identify DD participants, in other words, the precision was higher in ASD diagnosis compared to DD diagnosis independently of the dataset used. However, the confidence when the classifier marked a participant with DD was higher than the confidence when the classifier marked a participant with ASD, to put this in another way, recall was higher in DD diagnosis compared to ASD diagnosis for each dataset used. This resulted in higher F1 scores for DD diagnosis compared to ASD diagnosis.

**Table IV.** Machine Learning Results: the mean and the standard deviation of the measures obtained during 100 repetitions in the training and the testing of a Random Tree classifier for each dataset. (tn - true negative, fp - false positive, fn - false negative, tp - true positive, N features - the total number of features used to train the classifier, Acc train - accuracy of the classifier in the train set, Acc test - accuracy of the classifier in the test set, AUC - area under the roc curve, DD precision - precision measured for DD class, DD recall - recall of DD class, ASD precision - precision of ASD class, ASD recall - recall of ASD class, DD f1-score - F1-score of DD class, ASD f1 score - F1-score of ASD class. Positive class - ASD; Negative class - DD; Train size - 1846 participants (923 ASD / 923 DD); Test Size - 790 participants (395 ASD / 395 DD))

| Dataset |  | tn | fp | fn | tp | N features | Acc train (%) | Acc test (%) | AUC | DD precision (%) | DD recall (%) | ASD precision (%) | ASD recall (%) | DD f1 score (%) | ASD f1 score (%) |
| --- | --- | --- | --- | --- | --- | --- | --- | --- | --- | --- | --- | --- | --- | --- | --- |
| Gene Dosage | mean | 369.6 | 25.4 | 91.7 | 303.3 | 117.4 | 88.59 | 85.18 | 0.85 | 80.15 | 93.57 | 92.30 | 76.79 | 86.33 | 83.82 |
|  | sd | 5.3 | 5.3 | 7.7 | 7.7 | 3.9 | 0.41 | 1.11 | 0.01 | 1.35 | 1.35 | 1.47 | 1.96 | 0.98 | 1.31 |
| GO | mean | 368.0 | 27.0 | 105.5 | 289.5 | 62.0 | 86.25 | 83.22 | 0.83 | 77.73 | 93.15 | 91.50 | 73.28 | 84.73 | 81.36 |
|  | sd | 6.6 | 6.6 | 7.4 | 7.4 | 1.5 | 0.41 | 1.09 | 0.01 | 1.18 | 1.68 | 1.81 | 1.87 | 0.99 | 1.28 |
| Gene Dosage - GO | mean | 371.8 | 23.2 | 94.3 | 300.7 | 58.6 | 88.59 | 85.13 | 0.85 | 79.79 | 94.13 | 92.87 | 76.13 | 86.36 | 83.66 |
|  | sd | 5.0 | 5.0 | 7.2 | 7.2 | 1.6 | 0.37 | 1.06 | 0.01 | 1.24 | 1.27 | 1.42 | 1.83 | 0.94 | 1.25 |

Finally, we will focus on the number of features used to train and test the classifiers recorded in Table IV. On average, 117(sd 4) genes were needed to train a classifier with gene dosage data, whereas 62(sd 2) GO terms were used for the same purpose with a drop in the test accuracy of approximately ∼2%, in other words, with nearly half of the features was possible to train a classifier with almost the same performance by swapping the genes by its corresponding GO terms. This fact may impact the classifier ability of generalization (Bousquet, 2011; Bousquet & Elisseeff, 2002) in ML approaches applied in Autism due to the disorder low genetic occurrence and high genetic variability. For example, a classifier that relies only in gene dosage information to decide may not be able to give an accurate response in the presence of a gene that the classifier was not trained beforehand. On the other hand, if the classifier uses GO information it may be able to infer a correct response based on the previous examples because it is not tied to a set of genes but instead it is tied to a set of biological processes, molecular functions and cellular components relevant to the problem, indeed, a less variable and more compact set of features. However, we were aware that relying only in one type of data was not an optimal solution. As demonstrated earlier either genes or GO terms had different importance’s. In order to express the GO term importance in function of the gene importance we built the Gene Dosage - GO dataset. By using both gene and GO features we were able to reduce the required number of features to train the classifier to a minimum of 59 (sd 2) features without a drop on the classifier performance metrics.

## 4. Discussion

The present study has some unique features when compared to others addressing the same issue, both from the methodological and diagnostic points of view. In this discussion we will focus its sample size, accuracy obtained in ML diagnostic classification, ASD heterogeneity approach, and the value of gene and GO complex networks. In the end we will provide directions for developing based-knowledge systems addressing ASD classification (and in particular differential diagnosis) and future work on ASD ML approaches.

First, we used a sample size of 2.636 participants whereas in other studies aiming to diagnostic classification in ASD the maximum number of samples used was around 1000 (Abraham et al., 2017; Duffy & Als, 2012; Ghafouri-Fard et al., 2019; Ghiassian, Greiner, Jin, & Brown, 2016; Heinsfeld, Franco, Craddock, Buchweitz, & Meneguzzi, 2018; Parikh, Li, & He, 2019; Subbaraju, Suresh, Sundaram, & Narasimhan, 2017). Our best accuracy (85.18%) was only 2.02% behind the best classifier found in the studies above, which uses EEG brain features (Duffy & Als, 2012). Indeed, our study surpass in accuracy the study mentioned above in the genetic field by about 11.51% (Ghafouri-Fard et al., 2019).

Another characteristic of this work relates to its approach to ASD heterogeneity. We used a polygenic approach and map the participant relations with multiple genes encoding dopaminergic aspects. Furthermore, we go beyond gene duplication or deletion. Using GO terms, we explored and mapped the relations between participants and their genetic signatures, linking participants with similar disrupted biological process, molecular function s and cellular components within the dopaminergic pathway. Using complex network analysis, we demonstrated some unique characteristics of these maps and transform this information to address a differential diagnosis classification problem.

Given the results obtain in the present study, we consider viable its application for developing based-knowledge systems addressing support decision in ASD. One approach for developing support decision tools in this field should consider collecting and weighting the response of an ensemble of classifiers (Rokach, 2010) working with different types of data (ex.: genes, GO, brain) to deliver a diagnostic output response. Furthermore, to extend this approach beyond actions of the dopaminergic system in ASD, could be interesting to train several ML classifiers each one based on one type of system or pathway shown to be disrupted in ASD. It is likely that the performance of this approach depends on the type of differential diagnosis being performed. Combining the response of the classifiers will probably increase the diagnostic accuracy of the response while at the same time, analysing the individual output of each classifier may provide insights about the systems disrupted in a given ASD case.

We conclude that with our ML approach we could provide differential diagnosis between ASD and DD diagnosis using features extracted from CNV genes related to dopaminergic neurobiology aspects. However, there are other biologic agents, brain pathways and genetics contributing for ASD that could reveal new insights about the disorder using this approach. Thus, dopamine has impact on other neurobiological diseases that could not be studied here due to the small number of samples found in our sample. These other diseases and their dopaminergic characteristics should also be compared to the ASD dopaminergic characteristics. Furthermore, ASD is a neurodevelopmental disorder strongly tied to behavioural aspects. This is an opportunity to improve our knowledge about the brain and the genetic interplays involved in given neurotransmitter system. Future ML studies could explore this by building ML approaches to attempt predicting ASD psychometrics with neuroimaging and genetic features.

## Acknowledgements

We are grateful to all of the families at the participating Simons Simplex Collection (SSC) sites, as well as the principal investigators (A. Beaudet, R. Bernier, J. Constantino, E. Cook, E. Fombonne, D. Geschwind, R. Goin-Kochel, E. Hanson, D. Grice, A. Klin, D. Ledbetter, C. Lord, C. Martin, D. Martin, R. Maxim, J. Miles, O. Ousley, K. Pelphrey, B. Peterson, J. Piggot, C. Saulnier, M. State, W. Stone, J. Sutcliffe, C. Walsh, Z. Warren, E. Wijsman). We appreciate obtaining access to genetic data on SFARI Base. Approved researchers can obtain the SSC population dataset described in this study (https://gene.sfari.org/database/cnv/) by applying at https://base.sfari.org.

We thank the Portuguese Science Foundation (FCT) for financial support through “Projecto Operacional Regional do Centro”–BIGDATIMAGE: CENTRO-01-0145-FEDER-000016 and MEDPersyst POCI-01-0145-FEDER-016428, FCT, UID/NEU; UID/4950/2020, and the European Commission, H2020-SC1-2016-2017, STIPED

